# Structure of the native supercoiled flagellar hook as a universal joint

**DOI:** 10.1101/748384

**Authors:** Takayuki Kato, Fumiaki Makino, Tomoko Miyata, Peter Horváth, Keiichi Namba

## Abstract

Bacteria swim in viscous liquid environments by using the flagellum^1–3^. The flagellum is composed of about 30 different proteins and can be roughly divided into three parts: the basal body, the hook and the filament. The basal body acts as a rotary motor powered by ion motive force across the cytoplasmic membrane as well as a protein export apparatus to construct the axial structure of the flagellum. The filament is as a helical propeller, and it is a supercoiled form of a helical tubular assembly consisting of a few tens of thousands of flagellin molecules^4^. The hook is a relatively short axial segment working as a universal joint connecting the basal body and the filament for smooth transmission of motor torque to the filament^5,6^. The structure of hook has been studied by combining X-ray crystal structure of a core fragment of hook protein FlgE and electron cryomicroscopy (cryoEM) helical image analysis of the polyhook in the straight form and has given a deep insight into the universal joint mechanism^7^. However, the supercoiled structure of the hook was an approximate model based on the atomic model of the straight hook without its inner core domain^7^ and EM observations of supercoiled polyhook by freeze-dry and Pt/Pd shadow cast^8^. Here we report the native supercoiled hook structure at 3.1 Å resolution by cryoEM single particle image analysis of the polyhook. The atomic model built on the three-dimensional (3D) density map show the actual changes in subunit conformation and intersubunit interactions upon compression and extension of the 11 protofilaments that occur during their smoke ring-like rotation and allow the hook to function as a dynamic molecular universal joint with high bending flexibility and twisting rigidity.

---

We isolated polyhooks from a *fliK* deficient mutant strain of *Salmonella*, SJW880^9^, which produces polyhooks that are structurally identical to the native hook but grows as long as 1 µm^10^. We plunge froze a holey carbon grid loaded with this sample solution kept at a room temperature at least for a few hours before plunging into liquid ethane to keep the polyhooks in the native supercoiled form. The polyhooks were observed in motion corrected images (Fig. 1a), and about 35 nm long tubular segments were extracted with 90% overlap between the consecutive segments to obtain two-dimensional (2D) class averages, which clearly show helical arrays of FlgE subunits in different views (Fig. 1b). Then the 3D image was reconstructed from 157,334 such segment images (Fig. 1c). The resolution was 3.1 Å at a Fourier shell correlation of 0.143 (Fig. S. 1).

**Figure 1.**
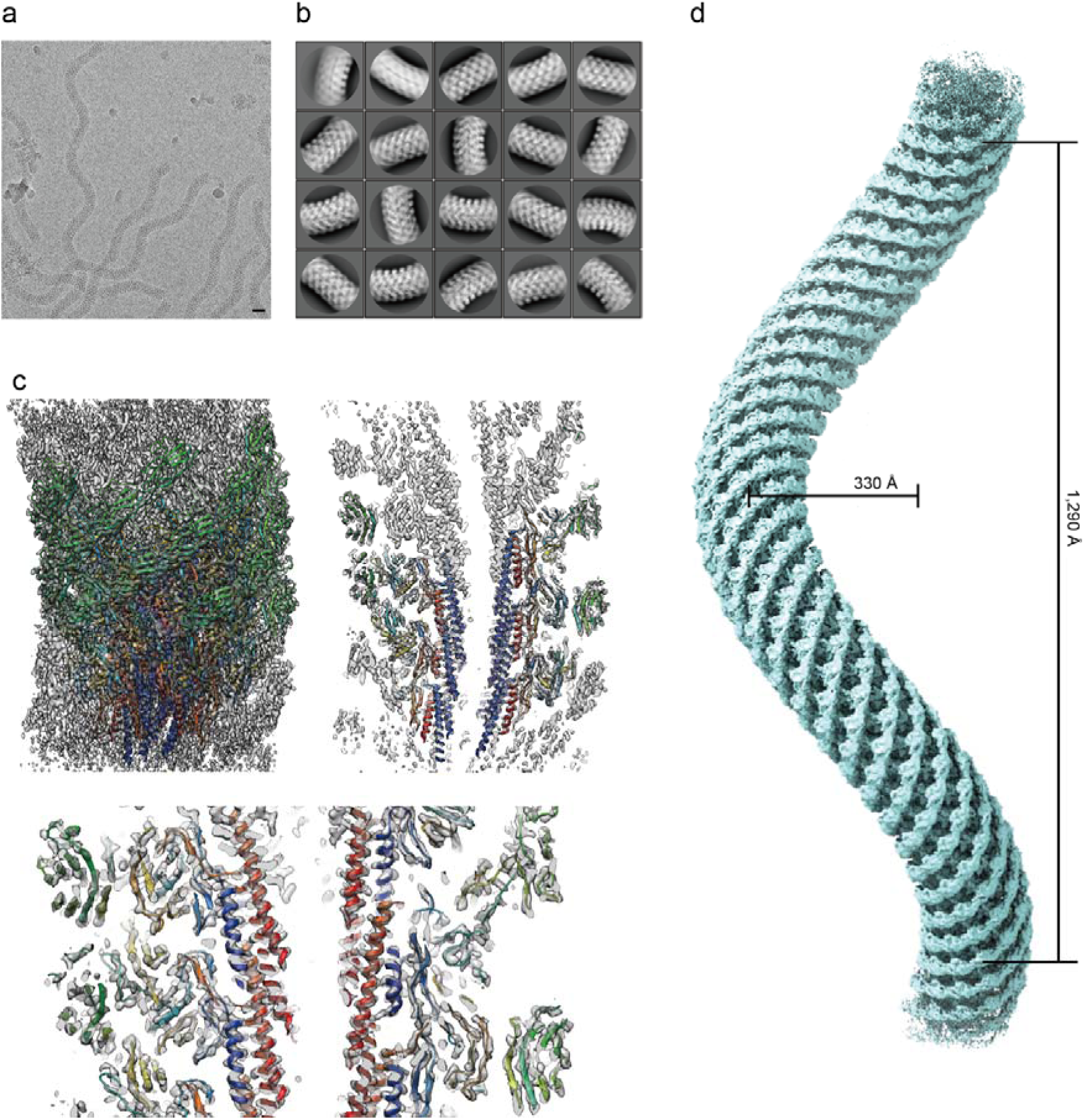
Structure of native supercoiled hook. **a,** A cryoEM image of ice embedded native supercoiled polyhook. **b**, Class averages by 2D classification. **c**, Reconstructed density map and atomic models of FlgE subunits in ribbon representation coloured in rainbow from the N- to C-terminus. **d**, A density map of supercoiled polyhook extended to the length of one helical pitch.

To estimate the helical pitch and radius of the supercoil, we built a long 3D density map of the polyhook by extending this short segment map to a length of one helical pitch of the supercoil (Fig. 1d, Movie S1). The supercoil was left-handed with a helical pitch of 129 nm and a diameter of 33 nm. Supercoiled forms of *Salmonella* polyhooks have been studied by EM with negative stain as well as freeze-dry and Pt/Pd shadow-cast to avoid flattening, under different solution conditions by changing pH and temperature^8^. The polyhook under neutral pH at room temperature was called “Normal”, and it was a right-handed supercoil with a helical pitch of 95 nm and a diameter of 35 nm. Somehow, the structure of the native polyhook we obtained is more similar to their “Left-handed” with a helical pitch of 60–100 nm and a diameter of 5 - 35 nm formed under pH 2 - 5 and at a temperature lower than 15°C, but no such large variations in the helical pitch and diameter are observed in our cryoEM images. So we believe the supercoil of the native hook is left-handed as is now shown in the near-atomic resolution structure revealed by cryoEM.

Based on this cryoEM map, we built atomic models of FlgE for 26 subunits that cover more than two turns of the 1-start helix of its helical assembly (Fig. 1c). The model of FlgE consists of three domains, D0, D1 and D2, arranged from the inner core to the outer surface of the hook, just as previously modeled^7,11^. But we were also able to build a model for residues 24 – 71 including domain Dc as a long β-hairpin (Ser 33 - Phe 60) and an extended chain (Thr 61 – Gly 71) formed by the region connecting the N-terminal α-helix of domain D0 and domain D1 as well as an extended chain connecting domain D1 to the C-terminal α-helix of domain D0 (Gly 358 – Asp 366) (Fig. S2).

The hook is made of 11 protofilaments^11,12^, just like the flagellar filament connecting to the distal end of the hook through two junction proteins^4^. Since the passes of the protofilaments are nearly parallel to the tubular axis of the hook, the protofilament length varies from one to the other. Accordingly, the subunit conformation gradually changes along the circumference from one protofilament to the next due to the supercoiled nature of the native hook structure, although the conformations of subunits are nearly identical within each protofilament. So, although the conformations of the 26 subunits we built on the map were all similar to one another, we identified 11 distinct conformations among them, one for each protofilament (Fig. S3). These 11 distinct conformations are all realized by slightly different arrangements of the three domains but the domain conformations are well preserved (Fig. S4). The color-coded distances between Cα atoms of corresponding residues of the axially neighboring subunits within each protofilament clearly visualize the curvature of the supercoiled hook (Fig. 2a). The short, middle and long distances are colored from cyan to dark pink, where the shortest distance was 36.9 Å in protofilament 7, and the longest one was 56.7 Å in protofilament 1 as numbered in the upper panel. These distances change as a function of the radius, and the average distances measured over Cα atoms in domains D2 were 38.4 and 53.6 Å, those at domains D1 were 40.6 and 50.4 Å, and those at domains D0 were 42.7 and 48.8 Å, respectively in protofilament 7 and 1 (Fig. S5). The subunit distance along the protofilament is 45.6 Å in the straight form of the hook^11^, which is close to the averages of these sets of paired two distances, indicating that the curvature of the native hook is formed by elastic bending deformation.

**Figure 2.**
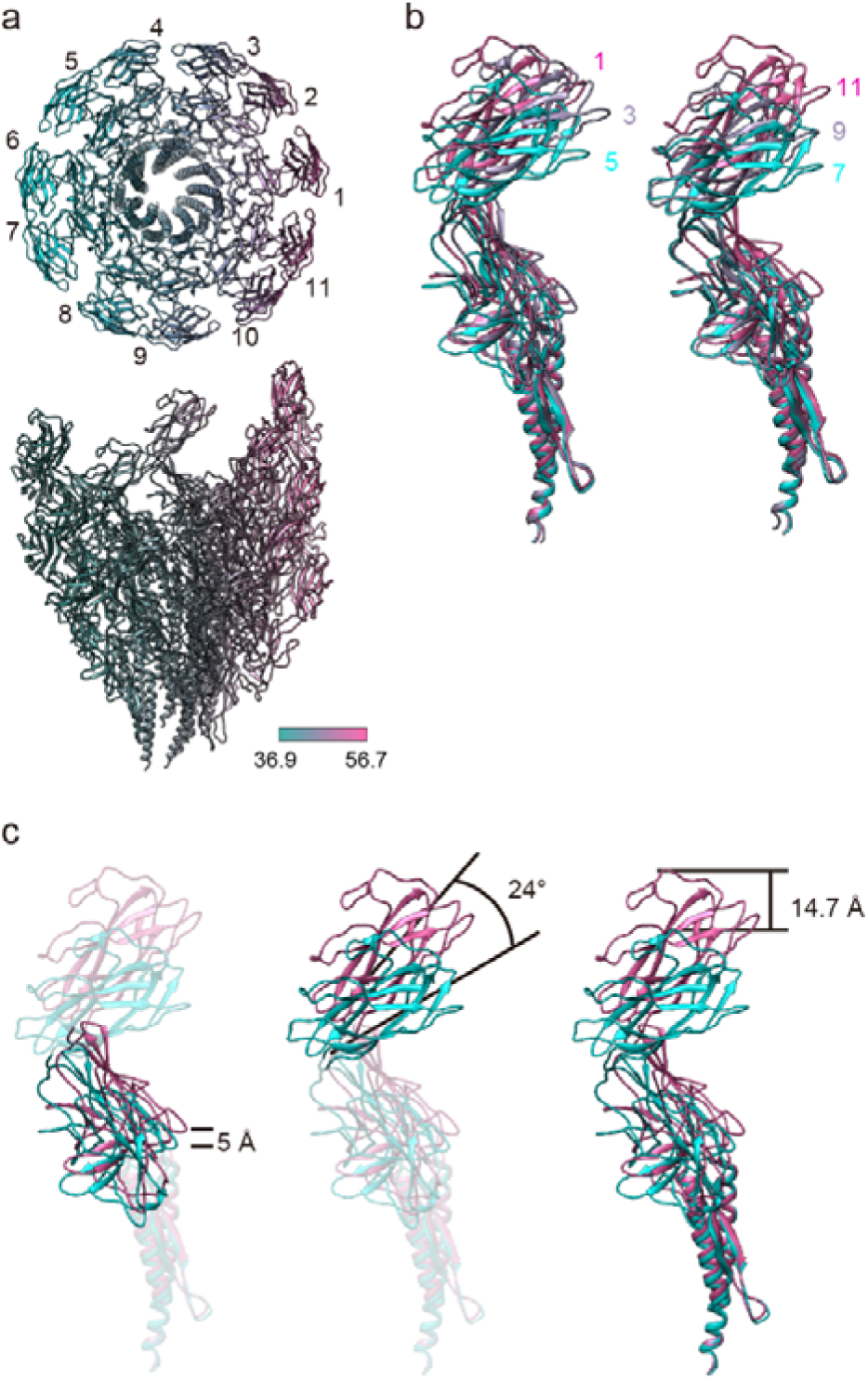
Conformational changes of FlgE subunits upon protofilament compression and extension. **a**, End-on and side views of the supercoiled hook colour-coded for the distance between corresponding residues of neighboring subunits along each protofilament. Each of the 11 protofilaments are numbered from 1 to 11, with protofilament 1 for the longest (extended) and 6 and 7 for the shortest (compressed). In end-on view, just one subunit is shown for each protofilament for clarity. **b**, Superimposition of FlgE subunits from protofilament 1, 3 and 5 (left) and from protofilament 7, 9 and 11 (right). **c,** Changes in position and orientation of domains D1 and D2 relative to D0 by its superposition. Colour codes are in a gradient from cyan for shortest (compressed) to pink for the longest (extended).

When the atomic models of FlgE in 11 different conformations are superimposed to each other, their gradual changes from one protofilament to the next are shown clearly (Fig. 2b, Movie S2). However, the superpositions of corresponding domains show almost no changes in their conformations as mentioned above (Fig. S4). When the two subunit conformations in the shortest and longest protofilaments are compared by superposing the D0 domains, domain D1 shifts up by 5 Å and tilts by 4°, and domain D2 tilts by 24° in addition, in order to achieve an overall subunit length extension of 14.7 Å (Fig. 2c). Thus, the compression and extension of each protofilament during the smoke-ring like rotation is achieved by changing relative domain orientations of FlgE while maintaining the conformation of each domain.

The smooth compression and extension of the protofilament are achieved also by well-designed intersubunit interactions. The overall changes of intersubunit interactions are depicted by comparing two sets of four subunit models, in which the middle two subunits are from the shortest and longest protofilaments, respectively (Fig. 3a). Domains D0 and Dc form one compact domain D0-Dc consisting of the N- and C-terminal α-helices and a long β-hairpin (residues 1-71 and 358-402), and this domain is tilted by about 17° from the tubular axis of the hook. Within each protofilament, the bottom of the N-terminal helix of the upper subunit is located on the top of the C-terminal helix with relatively large gaps of 6.3 Å and 12.5 Å in the short and long protofilaments, respectively (Fig. 3b). The bottom of the C-terminal helix of the upper subunit axially overlaps with the top of the C-terminal helix of the lower subunit to allow their mutual sliding for protofilament compression and extension (Fig. 3b and c). The relative disposition of the C-terminal helices between the neighboring protofilament is well maintained between subunits 0 and 5 but shows a slight axial shift between subunits 0 and 6 and a larger axial shift between subunits 0 and 11 (Fig. 3c). Residues Gly 329 – Asp 330 of subunit 0 at the tip of a short β-hairpin of domain D1 interact with residues Ala 39 – Asp 40 of subunit 5 in the distal part of the long β-hairpin of domain Dc, and this interaction does not change by protofilament compression and extension (Fig. 3d), indicating the importance of the long β-hairpin of domain Dc for the structural stability of the hook as found by deletion mutation experiments^13^.

**Figure 3.**
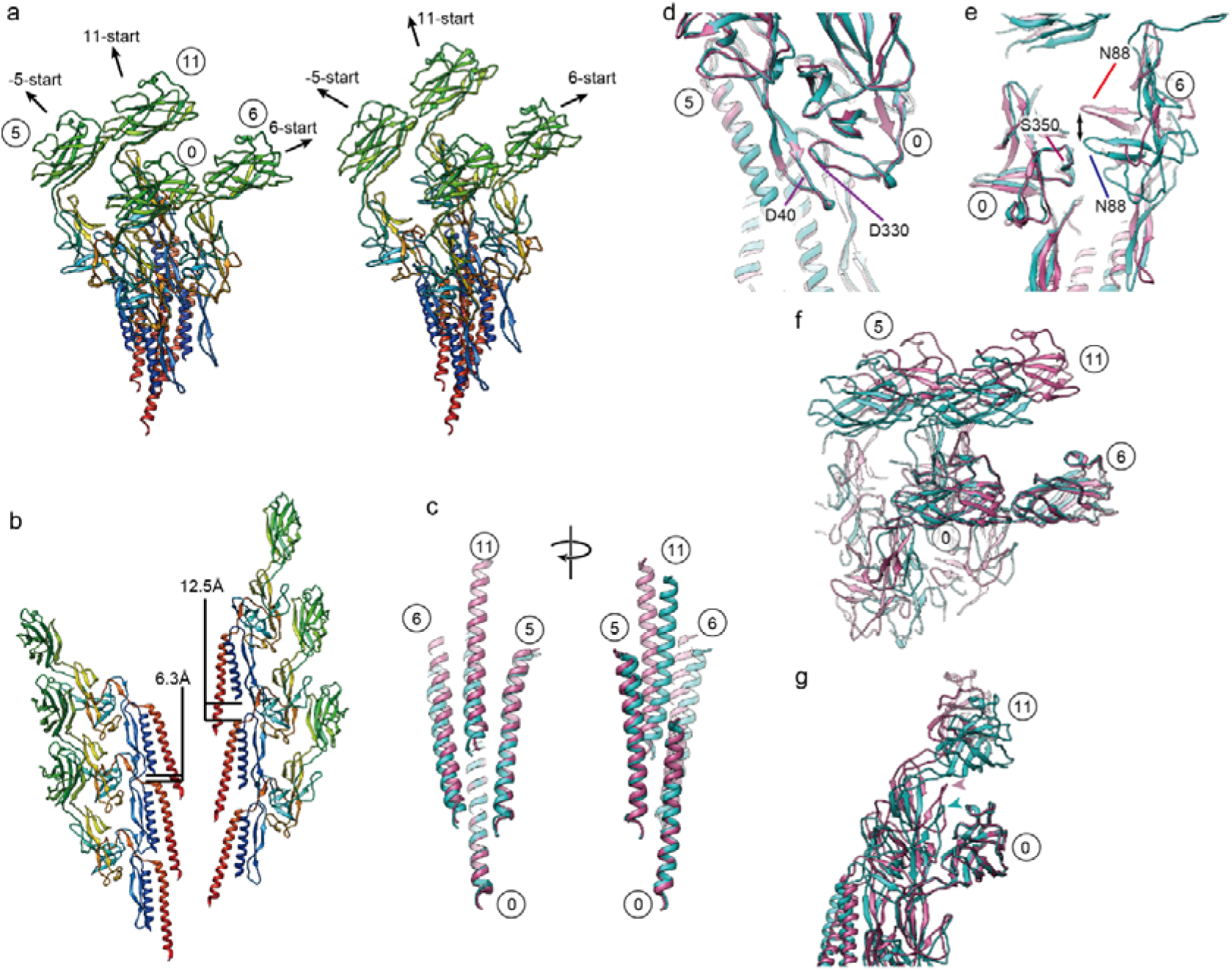
Changes in intersubunit interactions upon protofilament compression and extension. **a**, Conformations and intersubunit interactions of four neighboring subunits around the shortest (compressed) and longest (extended) protofilaments. A reference subunit is labeled 0 and the three neighboring subunits are labeled 5, 6 and 11 according to the three major helical directions of -5-, 6- and 11-start, respectively. **b**, The difference in the gap distance between the D0 domains along the protofilament for the shortest and longest. **c**, Superimposition of the C-terminal helices of subunits around the shortest and longest protofilaments. **d**, Constant interactions between domains D1 of subunit 0 and domain D0-Dc of subunit 5. **e**, Relative sliding shift between domain D1 of subunit 0 and domain D1 of subunit 6. **f**, Constant interactions between the D2 domains of subunits 0 and 6 along the 6-start helical directions on the surface of the hook. **g**, Relative sliding shift along the protofilament between domain D2 of subunit 0 and the triangular loop of domain D1 of subunit 11. The arrow heads indicate the triangular loop. In **a** and **b**, FlgE subunits in ribbon representation are coloured in rainbow from the N- to C-terminus, and in **c-g**, colour codes are in the same gradient as used in Fig. 2 from cyan to pink for the length of protofilament from the shortest to the longest.

The D1 domains form a mesh structure in the -5-, 6- and 11-start helical directions with some gaps^11^. These domain interactions stabilize the hook structure by maintaining the -5-start interactions while switching the 6-start interactions by an axial shift for protofilament compression and extension. Constant interactions are seen between residues Leu 101 – Glu 103 of subunit 0 with Ala 320 – Asn 321 and Gln 337 – Ser 339 of subunit 5 (Fig. 3d), but the interactions between Ser 87 – Asn 88 of subunit 6 and Gly 348 – Gly 350 of subunit 0 are present only in the compressed form (Fig. 3e cyan).

The D2 domains form a six-stranded helical array on the surface of the hook by their tight interactions along the 6-start helical direction (Fig. 3f) regardless of the protofilament length. The residues involved are Ser 159 – Thr 160, Asp 204 – Asn 205, Glu 233 – Asn 234 and Gln 268 – Thr 270 of subunit 0 with Leu 142 – Ala 145, Ser 188 – Asn 191, Asn 251 – Ala 253 and Lys 284 – Asp 287 of subunit 6. The close packing interactions between the D2 domains of the neighboring 6-start helical array on the inner side of the curved supercoiled hook were predicted to stabilize the native supercoiled hook structure^7^, and this was confirmed by a mutation experiment showing that the hook becomes straight by the deletion of domain D2 while keeping the bending flexibility to work as a universal joint^14^. We now see these interactions in the present structure (Fig. 1c). Domain D2 of subunit 0 maintains an axial intersubunit interaction along the protofilament with the triangular loop (Gly 117 – Pro 135) of domain D1 of subunit 11, and molecular dynamics simulation of the previous protofilament model showed a relatively large axial sliding upon compression and extension of the protofilament^7^. This is also confirmed in the present structure (Fig. 3g), indicating the importance of this dynamic intersubunit axial interaction for the bending flexibility of the hook while keeping its structural stability, as was also suggested by a mutation study^15^.

The stable interdomain interactions unperturbed by the protofilament compression and extension as well as variable interdomain dispositions that allow the hook to take supercoil conformations can be quantitatively measured from the distances between Cα atoms of corresponding residues from each of the 11 subunits in distinct conformations to their neighboring subunits in the -5, 6 and 11 major helical directions (Fig. 3a). The measured distances are depicted by color coding in Fig. S6. In the 11-start interactions (Fig. S6a), the tip of the C-terminal helix has the minimum standard deviation, and domain D2 shows the largest standard deviation, indicating that the bottom of the C-terminal helix is the pivot point for subunit tilt for hook bending. In the -5-start direction (Fig. S6b), domain D0 and the left half of domain D1 show small standard deviations. In the 6-start direction (Fig. S6c), the four-stranded β-sheet on the outside of domain D2 shows a smallest standard deviation because of their stable interactions along the 6-start helical line. Thus, the hook structure is composed of three radial layers formed by the D0-Dc, D1 and D2 domains, and the stable interdomain interactions in different helical directions in each of the three layers as well as the flexible connections between the layers are the basic design architecture of the hook to work as a molecular universal joint while maintaining the rigid tubular structure against twisting for transmitting motor torque to the filament (Fig. 4, Movie S3).

**Figure 4.**
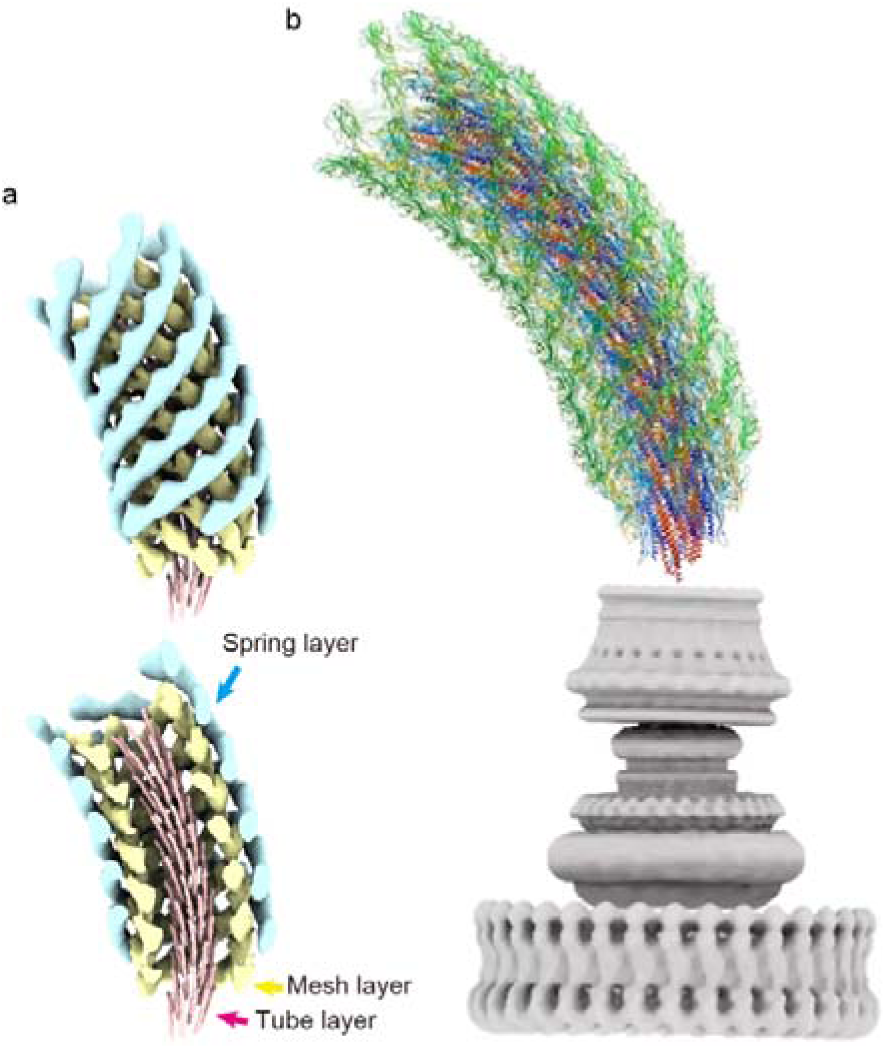
Three-layered architecture of the hook structure and the complete atomic model of the native hook as a universal joint. **a**, The tube, mesh and spring layers of the hook structure are colourd pink, yellow and cyan, respectively. **b**, The complete atomic model of the native hook with the flagellar basal body consisting of the C ring, MS ring and LP ring from the bottom to the top where the rod encapsulated inside the LP ring is directly connected to the bottom of the hook.

The rod protein FlgG shows considerable sequence and structural similarities to the D0, Dc and D1 domains of FlgE but does not have a domain corresponding to domain D2 of FlgE^16,17^. In contrast to the hook, however, the rod is straight and rigid to work as the drive shaft of the flagellar motor. This difference in the mechanical property results from the FlgG specific 18-residue insertion in domain Dc as was confirmed by an experiment in which the insertion of this sequence into FlgE made the hook straight and rigid^18^. FlgE of *Campylobacter jejuni* has a 17-residue insertion similar to this, and this insertion makes the β-hairpin of domain Dc much longer than that of *Salmonella* FlgE^19^. This β-hairpin was named L-stretch and occupies the gap formed by three D1 domains of subunits 0, 6 and 11^19^, possibly contributing to the higher stiffness of the *Campylobacter* hook. The corresponding gap in the *Salmonella* hook is much smaller in the compressed protofilaments (Fig. S7), suggesting that the insertion of the FlgG specific 18-residues forms the L-stretch just as that of *Campylobacter* hook to fill the gap to prevent protofilament compression in the hook made of the insertion mutant of FlgE. Therefore, this gap is essential for the bending flexibility of the hook.

Many different filamentous structures have been studied by cryoEM image analysis but the sample had to have a straight form with a helical symmetry by mutations or changing solution conditions. Now the cryoEM method has become powerful enough to solve even the structure of a native supercoil at near atomic resolution without any strict helical symmetry. So, this is another demonstration of the power of cryoEM technique in revealing physiologically meaningful macromolecular structures. The mechanism of polymorphic supercoiling of the flagellar filament to switch between left- and right-handed helical propellers for run and tumble of bacterial cell swimming for chemotaxis, which is distinct from that of the hook we presented here, is now within the reach.

## Acknowledgements

We thank Miki Kinoshita for analyzing the *flgE* sequence and Tohru Minamino for discussions and critically reading the manuscript. This work has been supported by JSPS KAKENHI Grant Number 25000013 to K.N. and 18K06155 to T.K and also supported by Platform Project for Supporting Drug Discovery and Life Science Research (BINDS) from AMED under Grant Number JP19am0101117 to K.N. and by JEOL YOKOGUSHI Research Alliance Laboratories of Osaka University to K.N.

## Author contributions

T.K. and K.N. conceived the project. T.M. and P.H. prepared the native hook sample. T. K. carried out data collection and structural analysis. T.K., F.M. and K.N. studied the structure and analyzed the conformational changes and intermolecular interactions of FlgE in the native hook structure. K.N. supervised this project. All authors discussed the results, and T.K., F.M. and K.N. wrote the manuscript on the basis of discussion with T.M. and P.H.

## Methods

### Sample preparation

A *Salmonella* polyhook strain, SJW880, was used to prepare polyhooks^20^ for cryoEM observation. The polyhook-basal bodies were purified in the same way as hook-basal body purification as previously described^21^. The polyhook-basal bodies were suspended in 25 mM Tris-HCl, pH 8.0, 100 mM NaCl. To prepare the native supercoiled polyhook, the purified sample was incubated overnight at room temperature.

### Data collection

A 2.4 μl sample solution was applied onto a glow discharged holey carbon grid (Quantifoil R1.2/1.3 Mo grid, Quantifoil Micro Tools, Germany), and the grid was plunge-frozen into liquid ethane by Vitrobot mark IV (Thermo Fisher Science, USA) with a blotting time of 3 second at 18 °C and 90 % humidity. All the data collection was performed on a prototype of CRYO ARM 200 (JEOL, Japan) equipped with a thermal field-emission electron gun operated at 200 kV, an Ω-type energy filter with a 20 eV slit width and a K2 Summit direct electron detector camera (Gatan, USA). An automated data acquisition program, JADAS (JEOL, Japan), was used to collect cryoEM image data, and pre-processing, motion correction and CTF estimation were carried out in real-time by the Gwatch image processing pipeline software we developed. Movie frames were recorded using the K2 Summit camera at a calibrated magnification of 45,579× corresponding to a pixel size of 1.097 Å with a defocus range from −0.6 to 1.8 μm. The data were collected with a total exposure of 10 s fractionated into 50 frames, with a total dose of ∼50 e^-^/Å^2^ in counting mode. A total of 1,702 movies were collected.

### Image processing

Motion correction was carried out by MotionCor2^22^, and the CTF parameters were estimated by Gctf^23^. Polyhook images were automatically selected by RELION 3.0^24^ as a helical object, and they were segmented into a box of 320 x 320 pixels with 90% overlap. After performing 2D classification for 1,029,196 such segmented images by RELION 3.0, the best particles were selected from the result. After ab-initio reconstruction from 323,254 segment images for 3 classes, 157,334 segment images of the best class were subjected to homogeneous refinement in cryoSPARC^25^. CTF refinement was carried out by RELION 3.0 using converted data from the cryoSPARC file format to the RELION file format. The final map was sharpened using a negative B-factor of 125 by cryoSPARC. Atomic model building was performed by COOT^26^, and then PHENIX^27^ was used for real-space refinement. All the images of the 3D maps and models used in this paper were prepared by UCSF Chimera^28^.

### Data availability

The cryoEM volume has been deposited in the Electron Microscopy Data Bank under accession code EMD-9952, and the atomic coordinates have been deposited in the Protein Data Bank under accession code 6K9Q.

