## Supplemental figure S1-S7 and a table S1 for "Structure of the native supercoiled flagellar hook as a universal joint"

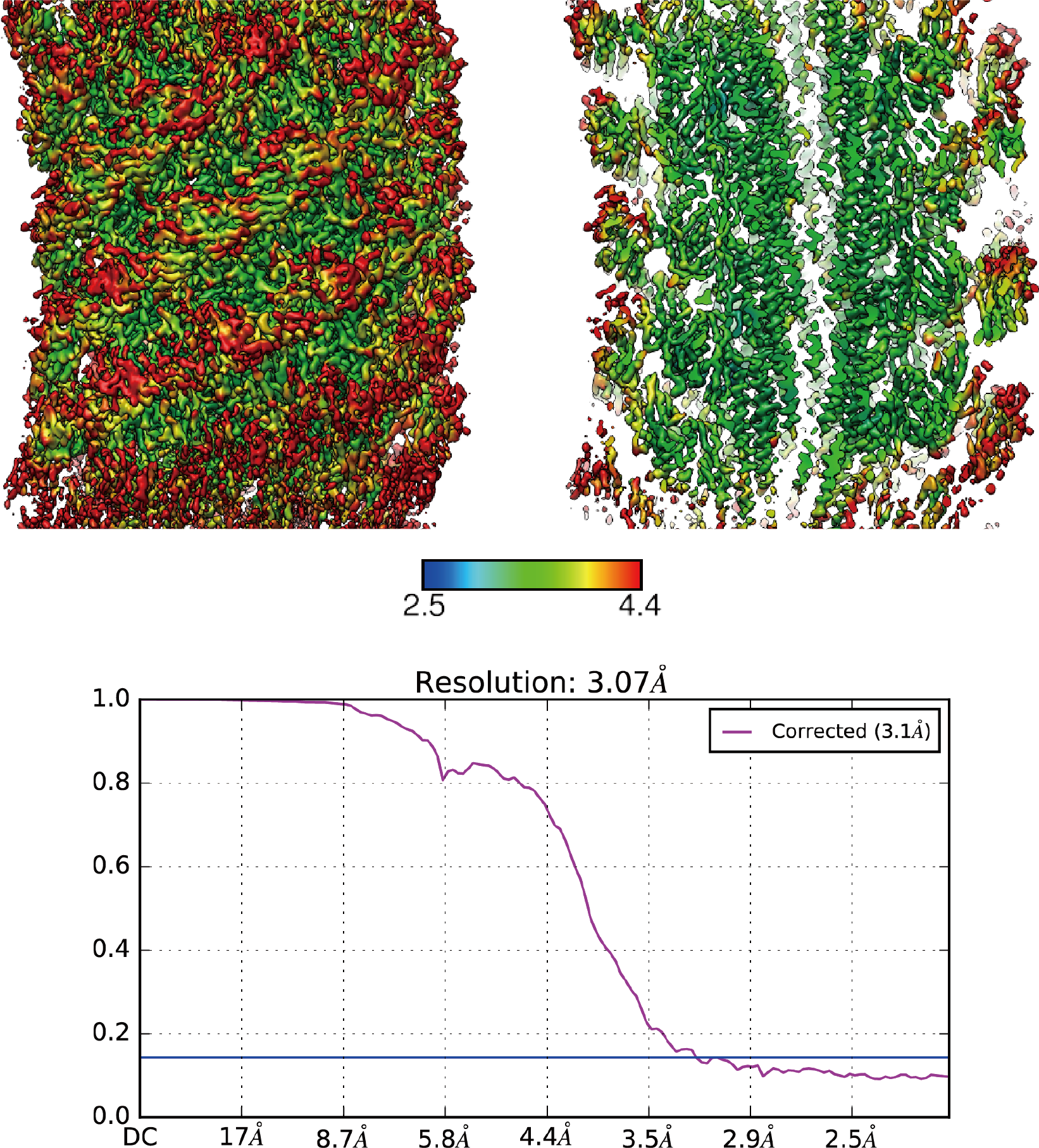

**Figure S1** Local and overall resolution of the native supercoiled hook structure. Colour maps of local resolution in the side view (upper left) and its cross-section (upper right). The blue to red gradient represents high to low resolution. The FSC curve of the reconstruction is shown in the lower panel.

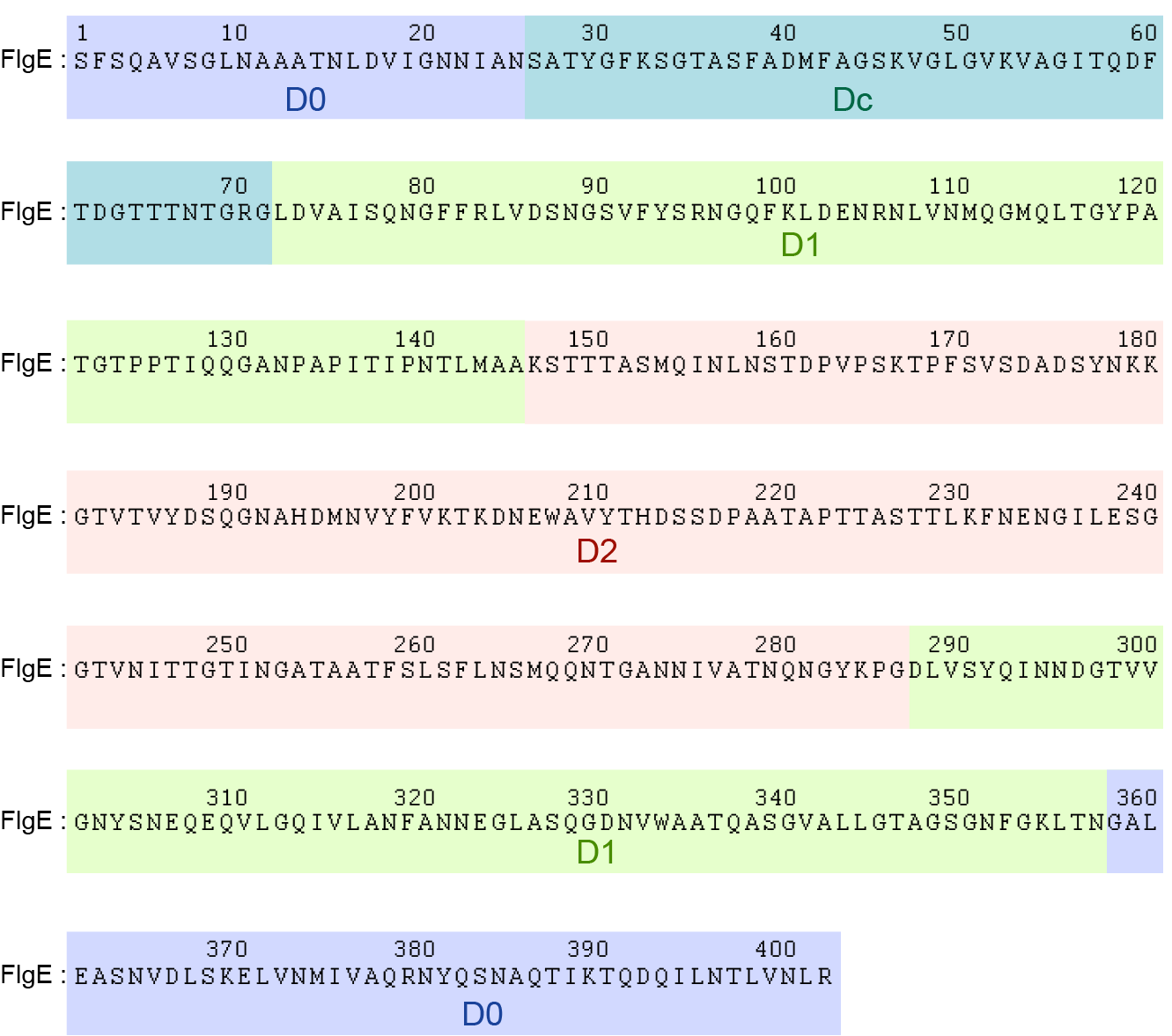

**Figure S2** Sequence of FlgE and regions of four domains, D0, Dc, D1 and D2. D0 (pale blue): Ser 1 – Gln 25, Asp 358 – Arg 402; Dc (blue-green): Ser 26 – Gly 71; D1 (light green): Thr 72 – Ala 145, Asp 286 – Val 365; D2 (pink): Lys 146 – Gly 286.

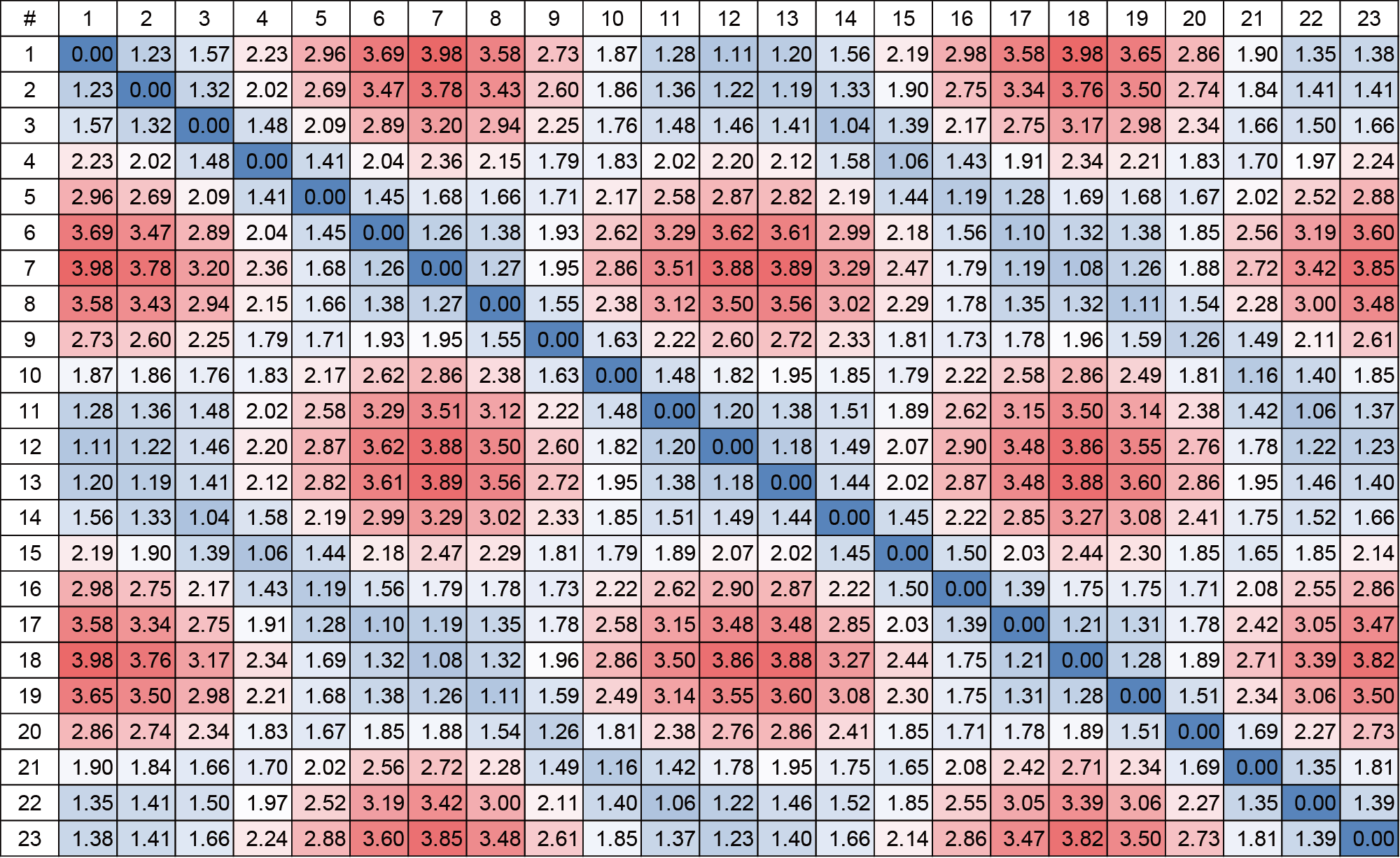

**Figure S3** Pairwise root mean square deviations (RMSDs) of Cα atoms between 23 atomic models of FlgE subunits in the supercoiled hook. Large and small RMSDs are coloured red and blue, respectively.

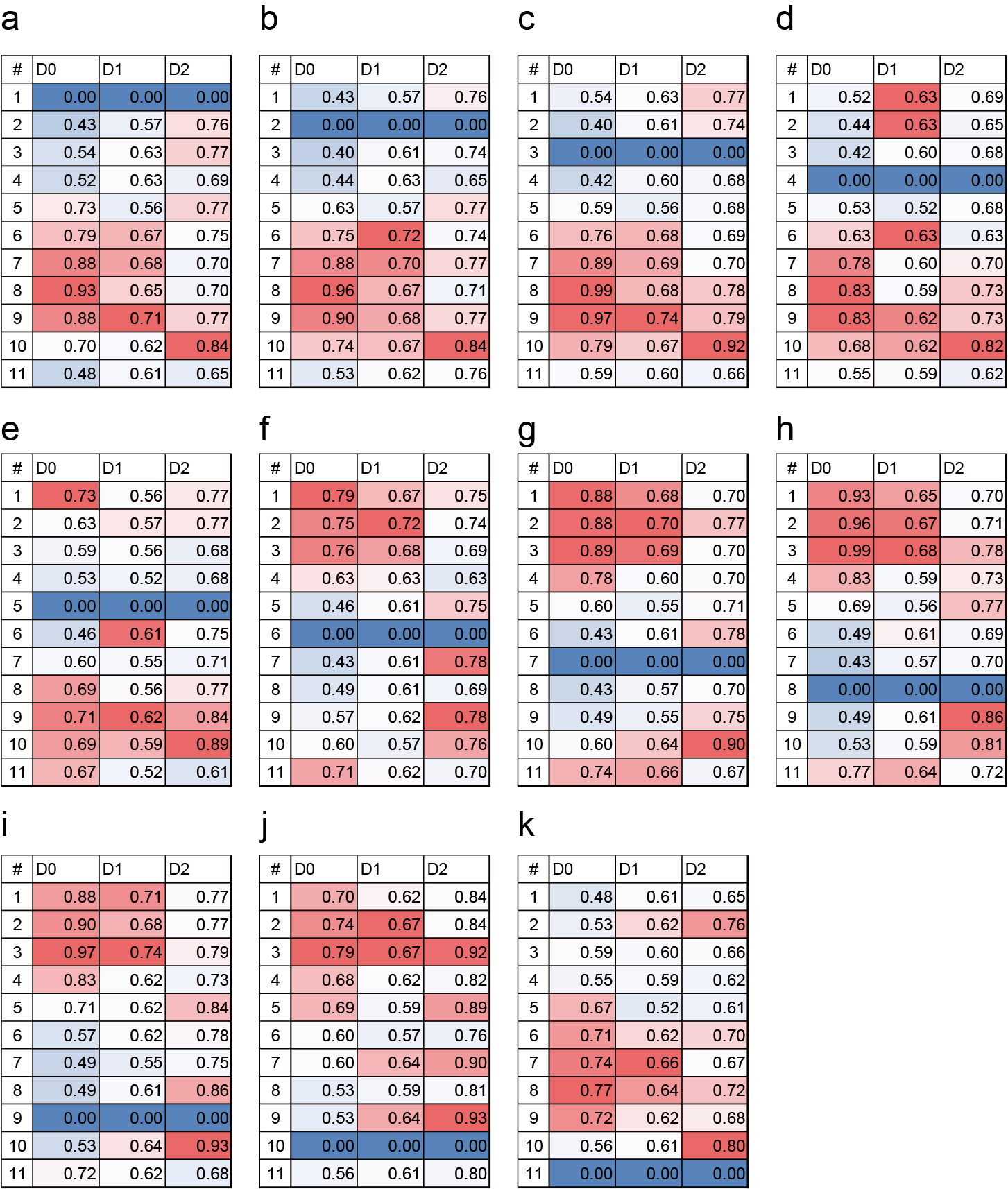

**Figure S4** Pairwise RMSDs of Cα atoms between 11 distinct conformations of FlgE subunit for each of three domains, D0, D1 and D2. Colours indicate the magnitude of RMSD with large and small RMSDs in red and blue, respectively. a – k, Each of 11 distinct conformations is used as a reference model for RMSD calculation.

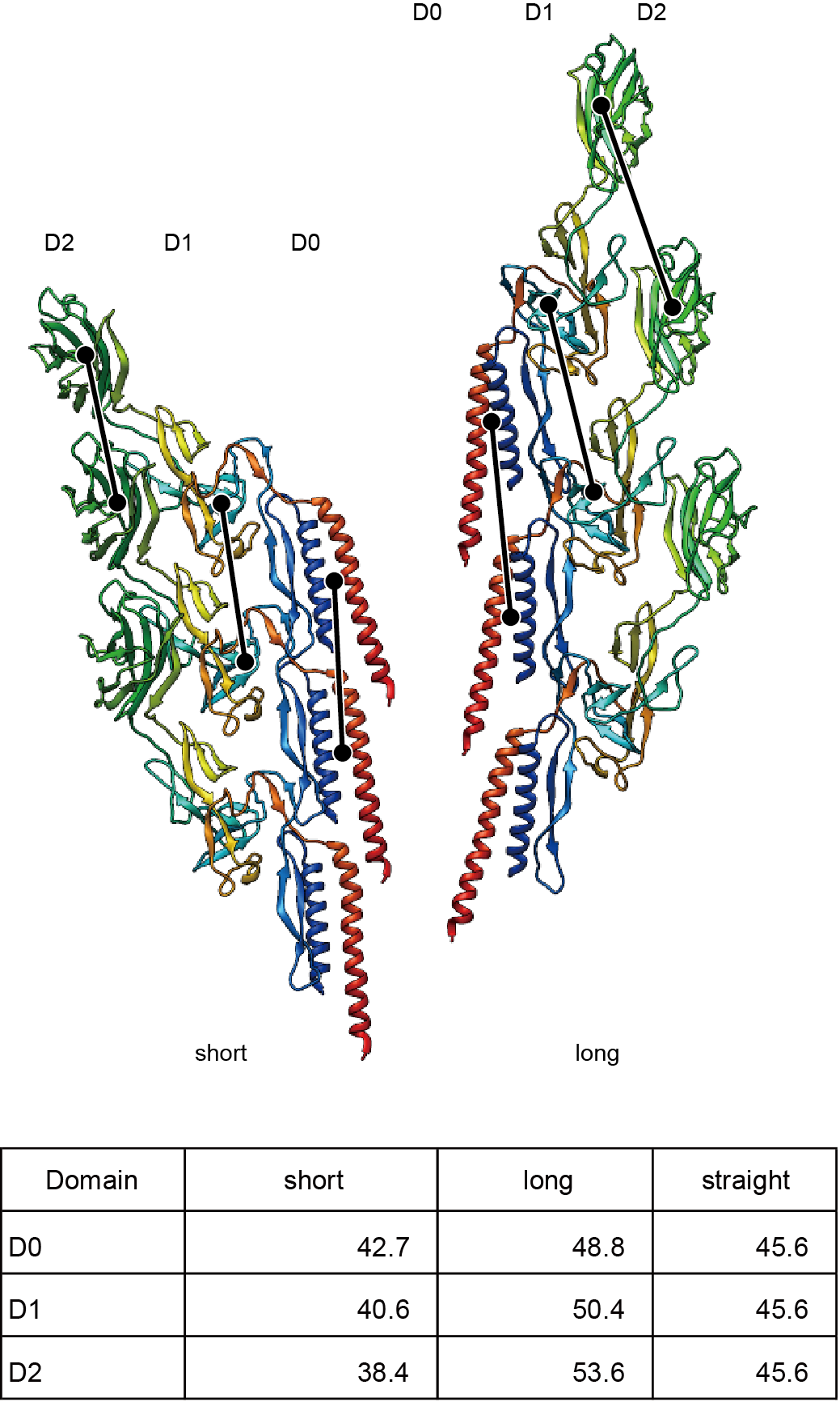

**Figure S5** Distances between neighboring subunits along the shortest and longest protofilaments and comparison with those of the straight hook structure. The distances are measured for each of three domains, D0, D1 and D2.Å

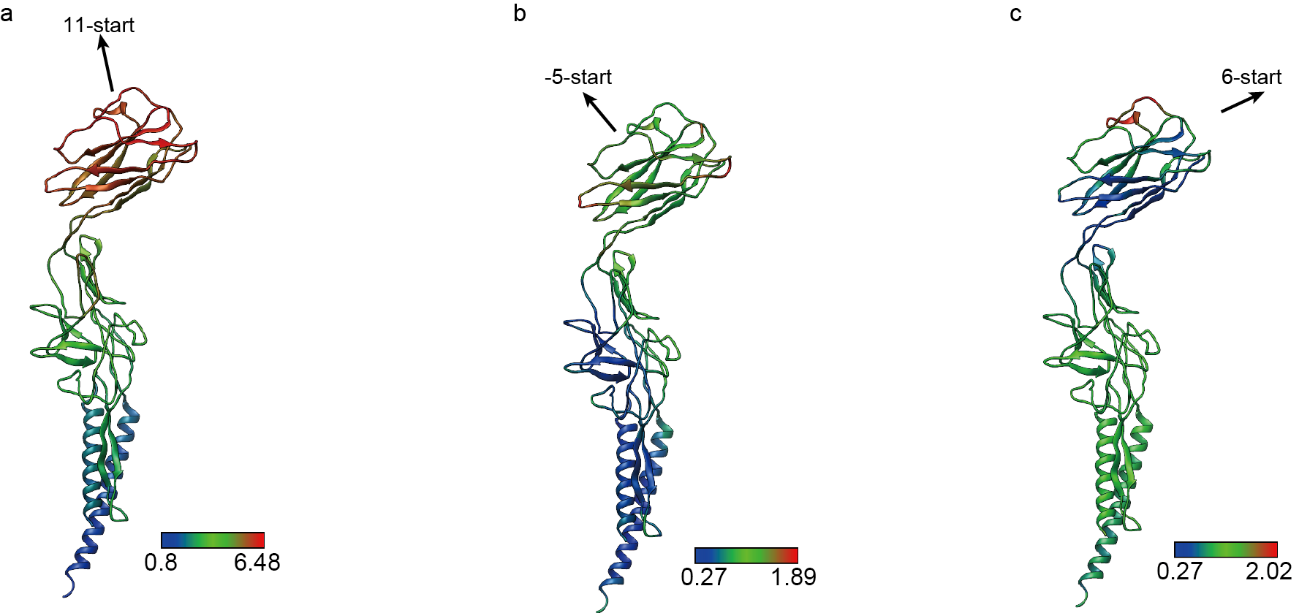

**Figure S6** Colour maps of standard deviations of distances between neighboring subunits in the 11-, -5 and 6-start helical directions measured over 11 distinct protofilament conformations. **a**, 11-start, **b**, -5-start, **c**, 6-start.

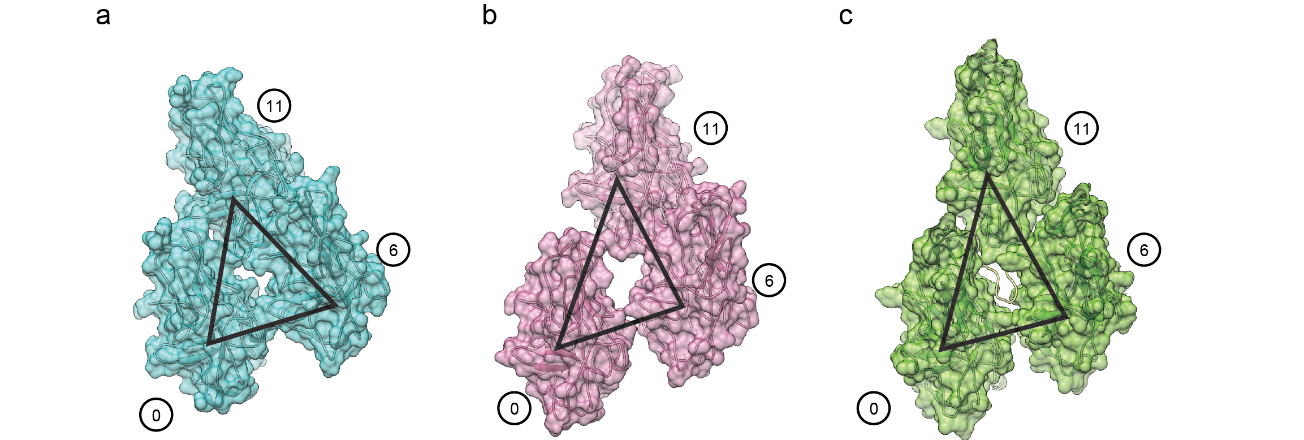

**Figure S7** The gap formed by three D1 domains of subunit 0, 6 and 11. **a**, Around the shortest protofilament, **b**, around the longest protofilament, **c**, for the straight hook of *Campylobacter jejuni*. In **c**, the loop in yellow filling the gap is the tip of the L-stretch of domain Dc of subunit 16.

**Table S1** Summary of cryoEM data collection, refinement and validation statistics

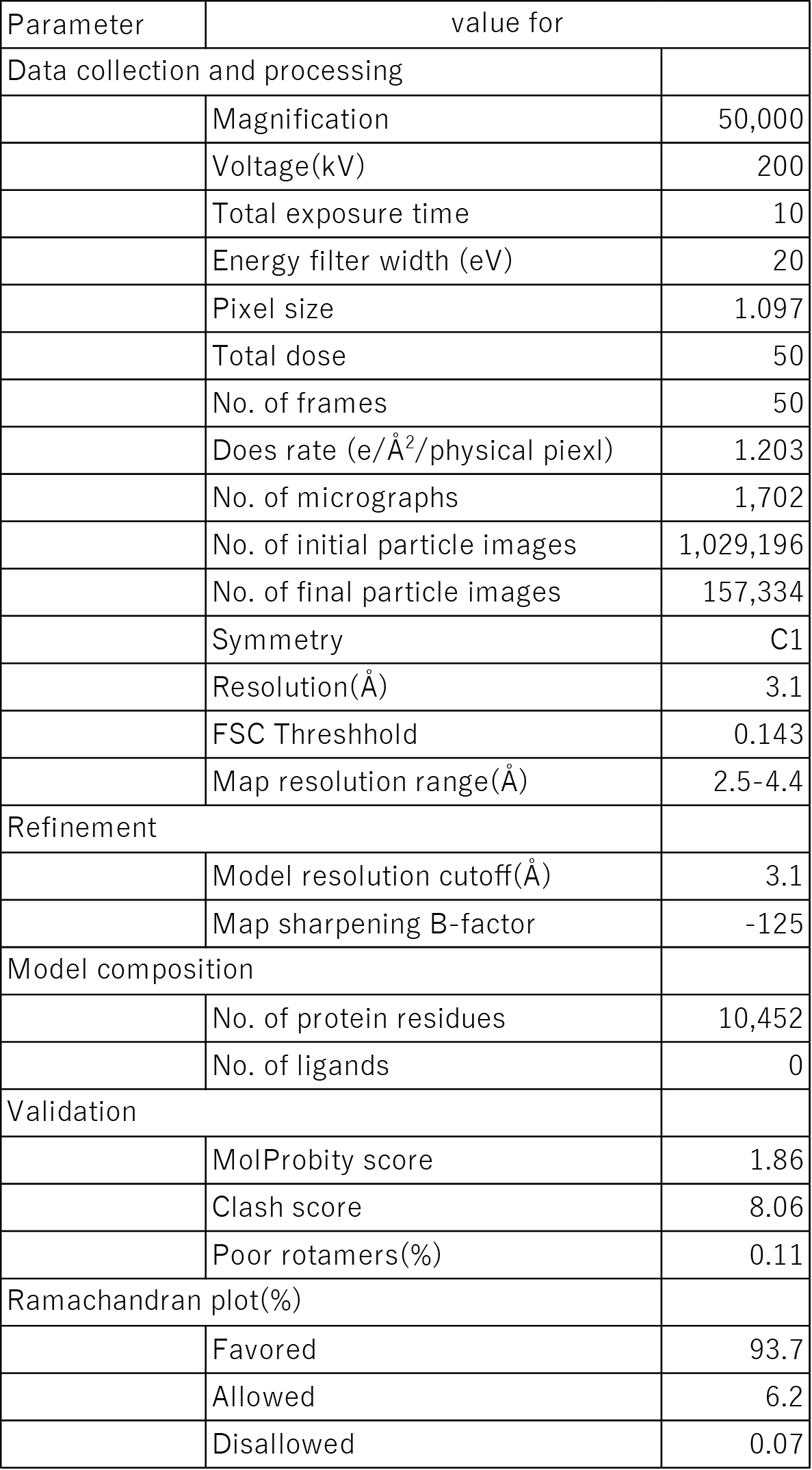
